# Contribution of Urea to Nitrite Production in Southern Ocean Waters with Contrasting Nitrifying Communities

**DOI:** 10.1101/2024.02.20.581251

**Authors:** J T. Hollibaugh, A. Okotie-Oyekan, J. Damashek, H. Ducklow, B. N. Popp, N. Wallsgrove, T. Allen

## Abstract

We compared the contribution of ammonia and urea to nitrite production in >100 samples of Southern Ocean waters with abundant and diverse ammonia-oxidizing archaeal (AOA) communities. Ammonia (AO) and urea (UO) oxidation rates were distributed uniformly within a water mass across coastal and slope waters west of the Antarctic Peninsula; however, rates and AOA community composition displayed strong vertical gradients. Rates in most samples from Antarctic surface and slope water were at or below the limit of detection. Highest mean rates of both processes were in the Winter Water (WW, epipelagic, 21.2 and 1.6 nmol N L^-1^ d^-1^), and the Circumpolar Deep Water (CDW, mesopelagic, 7.9 and 2.5 nmol N L^-1^ d^-1^), for AO and UO, respectively. However, we also found that the response of AO and UO to substrate amendments varied by water mass. AO rates in WW samples increased by ∼200% with 44 vs 6 nM amendments, but decreased (down to 7%) in CDW samples. UO rates responded similarly, but to a lesser degree. This response suggests that even low NH_4_^+^ amendments may inhibit AO by mesopelagic Thaumarchaeota populations. AO and UO rates were not correlated, nor were they correlated with the abundance or ratios of abundance of marker genes, or with the concentrations of ammonium or urea. Our data suggest that while ammonium is the primary substrate, urea-N is responsible for a significant fraction (∼25% of that from AO alone) of nitrite production in the Southern Ocean, comparable to its contribution at lower latitudes.

**IMPORTANCE:** Southern Ocean nitrification fuels denitrification in oxygen depleted zones at higher latitudes, one of the controls of N:P ratios in the global ocean. N_2_O, a powerful greenhouse gas, is by-product of nitrification. We contrast the contributions of ammonium and urea-N to nitrification in the Southern Ocean. Our work constrains rates and demonstrates that the contribution of urea-N to nitrite production in polar waters is comparable to that in temperate oceans. Correlations between activity and the abundance or ratios of Thaumarchaeota marker genes were weak, questioning their use as indicators of activity. We document differential responses of activity to substrate amendments by water mass: enhanced in epipelagic but inhibited in mesopelagic samples. We interpret this difference in the context of community composition and the production of reactive oxygen species. Our insights into environmental controls of nitrification are relevant to microbial ecologists studying Thaumarchaeota and to modeling the global nitrogen cycle.

## INTRODUCTION

Ammonia-oxidizing Thaumarchaeota (also referred to as Ammonia-Oxidizing Archaea, AOA, or the class *Nitrososphaeria*) play an important role in the nitrogen cycle by oxidizing ammonia to nitrite (1-3), and they are abundant in Antarctic coastal waters (4-6). Identification of genes for putative ureases and urea transporters in Thaumarchaeota genomes (7, 8) suggested that they might also be able to oxidize N supplied as urea, potentially increasing the rate of nitrite production *in situ* over that measured using ammonium. Subsequent work (9-11) demonstrated that the ability to oxidize urea-N is not universal in Thaumarchaeota, even among closely related isolates from the same environment. Alonso-Sáez et al. (12) used ratios of the abundance of Thaumarchaeota *ureC* to 16S rRNA (*rrs* hereinafter) or *amoA* genes, and incorporation into biomass of C supplied as urea, to infer that urea might be particularly important as a source of reduced N to Thaumarchaeota populations in polar (Arctic and Antarctic) waters. Results of their initial gene survey were replicated in subsequent work in the Arctic (13). Relatively few studies, and neither of these, have used ^15^N tracers to compare the oxidation rates of N supplied as urea (UO) and ammonium (AO) directly in the same sample. Recent work (14-17) has demonstrated that the contribution of urea to nitrification in the open ocean can be significant, if highly variable.

There are few measurements of UO in samples from Antarctic waters, thus the contribution of urea-N to nitrite production there, relative to AO or other processes, is understudied and poorly constrained. Pilot experiments performed using samples of Antarctic coastal waters found that the mean ratio of UO/AO in 3 samples from the Winter Water water mass was 1.9, while it was 0.3 in 3 samples from the Circumpolar Deep Water. A 2018 cruise to the continental shelf and slope west of the Antarctic Peninsula provided an opportunity to compare the relative contributions of urea-N and NH_4_^+^ to nitrite production in a larger data set and to perform process studies. We examined the response of AO and UO to substrate amendments to gain insight into the factors controlling rates *in situ* and to evaluate the effect of tracer additions on measured rates. We assessed the effect of incubation temperature on rates to evaluate the significance to rate measurements of deviations of incubation temperatures from *in situ*, and to assess the potential response of polar nitrification to warming oceans. We examined the correspondence between AO and UO rates and genetic markers for these processes to evaluate the use of gene ratios (12) as proxies for activity. Finally, we compared the relative contributions of UO and AO to nitrite production in samples from the Southern Ocean with their contributions at other locations.

## RESULTS

### Description of the study area

Cruise LMG1801 spanned 4 weeks during the Antarctic summer (6 January to 4 February, 2018, Supplemental Table 1) and sampled stations on the PAL LTER sampling grid, a strip of the continental shelf and slope west of the Antarctic Peninsula 700 km parallel to the coast by 200 km perpendicular to the coast (Supplemental Figure 1). This is a physically dynamic coastal ocean (18) in a region of extreme seasonality. There are 4 water masses in the study area (18, 19): Antarctic Surface Water (ASW, sampled at 10 or 15 m); the Winter Water (WW, sampled at the water column temperature minimum, 35-100 m depending on location); the Circumpolar Deep Water (CDW, sampled at 175 – 1,000 m); and Slope water (SLOPE, sampled at 2,500 to 3,048 m, generally ∼10 m above the bottom at stations on the slope or over basins on the shelf).

### Response of AO and UO to ^15^N amendments

Responses of WW versus CDW populations to ^15^N amendments differed markedly, as shown in Figure 1 and Supplemental Figure 2. Figure 1 plots rates measured at higher amendments normalized to rates measured with 6 nM amendments in the same sample as ((rate at [X]/rate at 6 nM)*100), versus substrate enrichment as (((amendment+ambient)/ambient)*100). This calculation assumes that 6 nM represents a true tracer addition with no effect on *in situ* rates, which is not necessarily correct. AO rates in WW samples increased (to >200%) with increasing amendments of ^15^NH_4_^+^. In contrast, AO rates in CDW samples were reduced significantly (to 7%) by increasing ^15^NH_4_^+^ amendments (Figure 1, Supplemental Table 2). This figure also shows that rates in both WW and CDW samples responded significantly to substrate enrichments that were <200%. UO rates in WW samples also increased with increasing ^15^N-urea amendments (Figure 1, Supplemental Table 2), while UO rates in CDW samples decreased with increasing urea amendments. CDW populations had a stronger response to NH_4_^+^ than to urea amendments (Figure 1); however, the difference was not significant (2-tail *t*-tests, *p*=0.069 and *p*=0.081 for 44 vs 6 and 440 vs 6 nM amendments, respectively).

**Figure 1.**
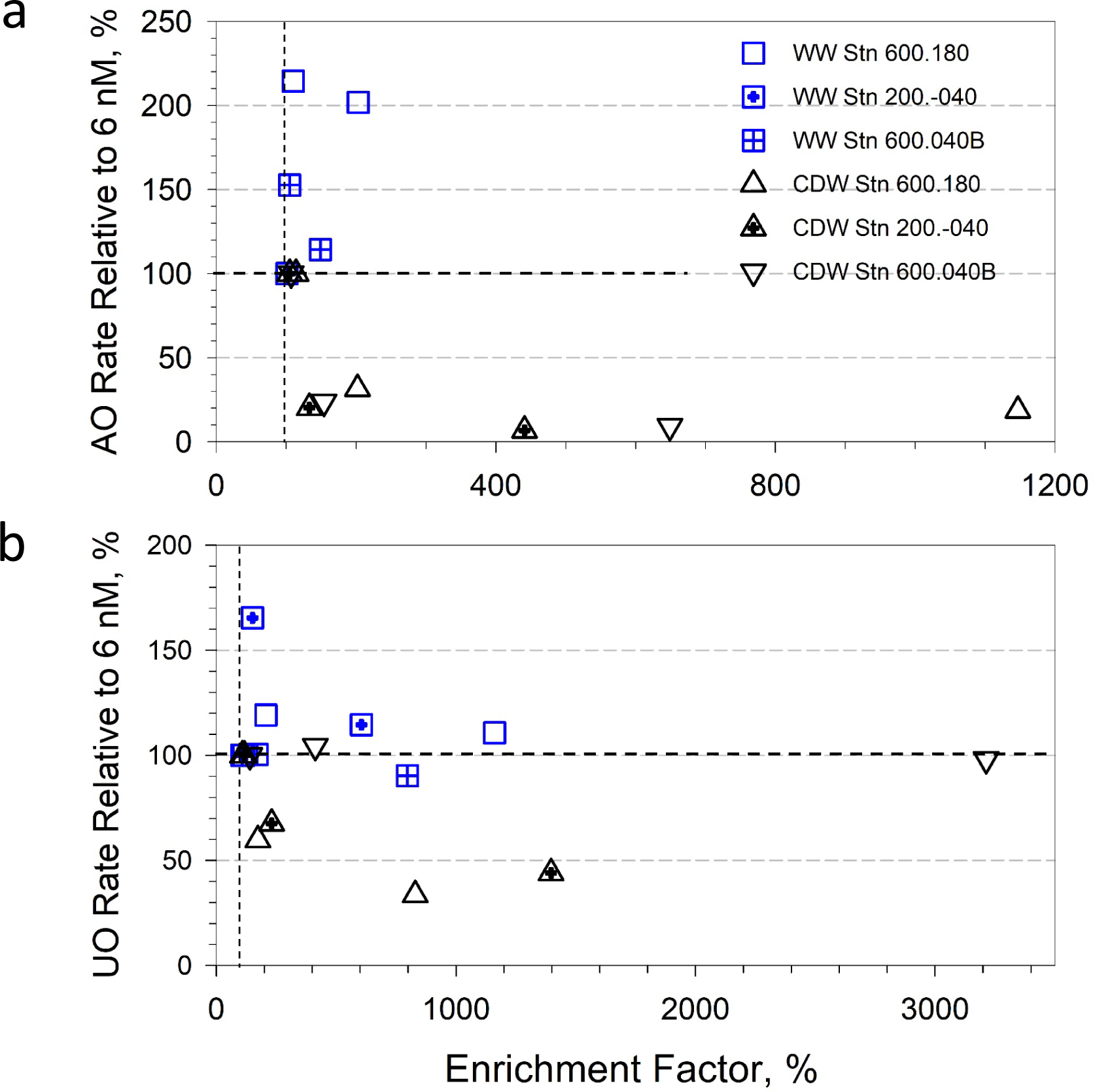
Oxidation rates of N supplied as NH_4_^+^ (AO, panel a) or urea (UO, panel b) as functions of amendments with ^15^N-labeled substrates (as nmol L^-1^ of the substrate, not of N in the case of urea). This figure combines data from 3 stations as indicated by the legend in panel a. Sample depths are given in Supplemental Table 1. Rates have been normalized relative to the rate measured in the same sample with a 6 nM amendment: ((rate at X nM/rate at 6 nM)*100). Amendments are normalized (as “enrichment factor”) as the increase in total substrate concentration relative to ambient: (((amendment + ambient)/ambient)*100). Vertical and horizontal dashed lines show 100%: no enrichment and no response, respectively.

### Response of AO to incubation temperature

Production of ^15^NO_x_ from ^15^NH_4_^+^ (we did not test urea) increased with temperature to maxima at 5-10 °C, then declined. The same pattern was seen with samples from two different stations and with both WW and CDW (Supplemental Figure 3). We found that rates were greater than the limit of detection (>LD) in incubations at 0 °C at all stations and depths tested, and were >LD in incubations at -1.0 °C in 3 of the 4 samples tested. Mean Q_10_ values for AO calculated for the interval 0 to 3 or 5 °C averaged 2.24; (Supplemental Table 3), similar to the value (1.1) reported by (20). However, Q_10_ values calculated for the interval -1.8-0 °C were much larger: 12.3-14.7. The Percival^®^ incubator we used maintained sample temperatures at (median, max, min) 0.25, 2.85, -1.50 °C, while *in situ* temperatures for our samples were: WW, -1.28, 0.16, -1.69; and CDW, 1.40, 2.04, -0.04 (Supplemental Table 4). The medians of AO rates measured in WW and CDW samples are 9.1 and 5.1 nmol N L^-1^ d^-1^. Assuming Q_10_=2.24 applies to all of our samples, medians of AO rates *in situ* would be 8.0 and 5.6 nmol L^-1^ d^-1^, or 0.89 and 1.1 times the rates we report. A similar calculation using the mean Q_10_ for the interval -1.8 – 0 °C (13.5) yields a median *in situ* rate for WW samples of 6.1 nmol L^-1^ d^-1^, or ∼70% of the measured rate. We assume these corrections would apply to UO rates as well. We have not corrected the data reported in Supplemental Table 1 for the 10 to 30 % error due to differences between *in situ* and incubation temperatures. This observation also suggests a relatively small change in nitrite production by AO, driven strictly by temperature, in a warming Southern Ocean.

### Variation within water masses

The study area has a strong seasonal cycle and complex physical oceanography tied, in part, to melting ice. We examined data from the WW and CDW water masses to determine if they displayed a temporal signal by splitting the data set into two groups representing samples collected at the beginning (days 1-15, n=104) versus end (days 16-29, n=60) of the cruise. Mann-Whitney ranks tests of the null hypothesis that values were distributed uniformly between these two groups revealed that UO rate was the only variable with a significant (*p*<0.05) temporal signal (Supplemental Table 5). UO rates were higher (8.4 vs 1.2 nmol N L^-1^ d^-1^) in CDW samples collected near the beginning of the cruise.

We used the same approach to determine if there were gradients within a water mass in the distributions of variables across the study area (Supplemental Figure 4). We restricted our analysis to WW and CDW water masses as many of the values for some variables were <LD in samples from the ASW and SLOPE water masses. We grouped samples by station location (northeast, n=88 versus southwest, n=76; and inshore, n=86 versus offshore, n=78), as shown in Supplemental Figure 1. Median AO rate was significantly higher in CDW samples from the NE end of the sampling grid (10.4 vs 3.2 nmol N L^-1^ d^-1^, *p*<0.05) and at inshore stations (9.0 vs 4.9 nmol L^-1^ d^-1^, *p*<0.05). Median UO rates were greater in WW and CDW samples from stations on the NE end of the sampling grid (2.0 vs 0.8 nmol N L^-1^ d^-1^ and 8.8 vs 1.4 nmol L^-1^ d^-1^, respectively; *p*<0.05; Supplemental Table 5). WW samples were both warmer and saltier at the NE end of the sampling grid, while CDW samples were warmer at offshore stations (Supplemental Table 5).

### Differences between water masses

Rates of AO and UO differed significantly (*p*=0.008) between water masses (Figure 2, Supplemental Table 6). Rates measured in samples from the WW averaged 21.2 and 1.6 nmol N L^-1^ d^-1^ (values <LD set to 0, (21), while those in CDW samples averaged 7.9 and 2.5 nmol N L^-1^ d^-1^ for AO and UO, respectively (Supplemental Table 4). AO and UO rates were <LD (<4.3 and <0.6 nmol N L^-1^ d^-1^ for AO and UO, respectively) in many of the samples from the ASW and SLOPE water masses (Supplemental Table 1). Means over all samples of the oxidation rates of N supplied as NH_4_^+^ or urea were 10.8 (n=216, range 0-158) and 2.5 (n=217, range 0-120) nmol N L^-1^ d^-1^, respectively (Supplemental Table 4). The highest UO rates (114 and 120 nmol N L^-1^ d^-1^, Figure 2b) were from replicates of one CDW sample with an elevated urea concentration (2,060 nM). If these outliers are removed, the mean UO rate is 1.5 nmol N L^-1^ d^-1^ (range 0-14).

**Figure 2.**
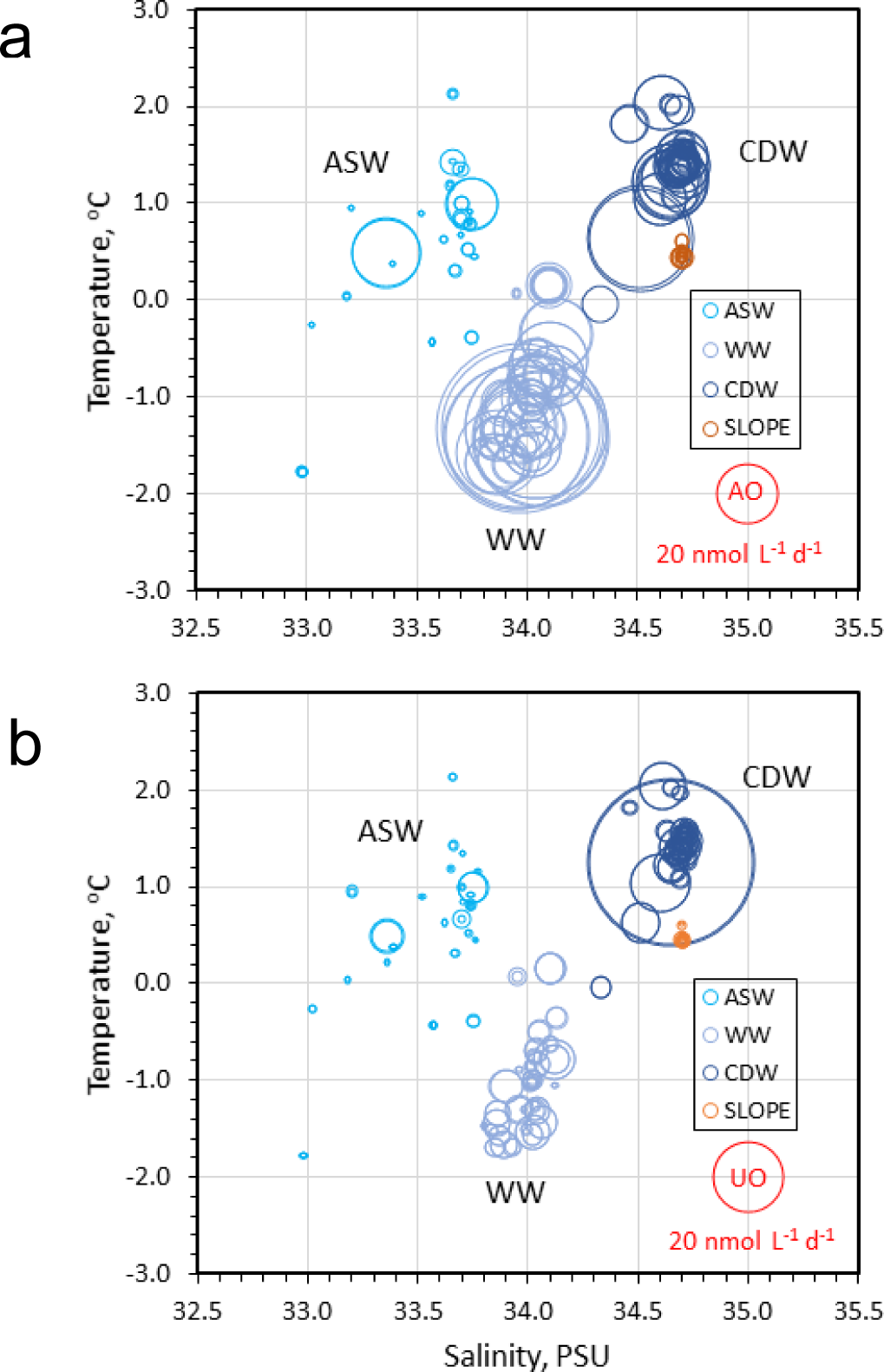
Rates of AO (panel a) and UO (panel b) plotted against *in situ* temperature and salinity. The origins of samples by target water mass are indicated by line colors. Samples with no activity have been assigned values of 0.1 nmol L^-1^ N d^-1^. The areas of the red bubbles at (35, - 2) are scaled to 20 nmol N L^-1^ d^-1^

Ammonium concentrations were greatest in samples from the ASW and WW, with mean concentrations of 930 and 620 nM that were not significantly different at *p*<0.01 (Supplemental Table 4, Supplemental Table 6). Ammonium concentration decreased with depth to mean concentrations of 160 and 200 nM in CDW and SLOPE samples, respectively. Ammonium concentrations were generally higher than those of urea (averages of 500 versus 130 nM over all samples), with no statistically significant differences between water masses (Supplemental Table 4, Supplemental Table 6). Both data sets contained outliers that were excluded from these calculations and NH_4_^+^ data are missing for some samples. The mean ratios of N available as urea versus NH_4_^+^ were 0.34, 0.36, 0.95 and 0.31 in ASW, WW, CDW and SLOPE water samples, respectively, if one outlier from a SLOPE water sample (urea concentration 1,800 nM, resulting in a urea-N/NH_4_^+^ ratio of 57) is excluded.

The abundances of all of the genes we measured (Supplemental Table 4, Supplemental Figure 5) were statistically significantly different (*p*<0.01) between the 4 water masses (Supplemental Table 6). The mean abundance of 16S rRNA (*rrs* hereinafter) from Bacteria decreased with increasing depth from 1.3 x 10^9^ copies L^-1^ in samples from ASW to 0.01 x 10^9^ copies L^-1^ in samples from SLOPE water. In contrast, the mean abundance of Thaumarchaeota *rrs* increased from 550 x 10^3^ copies L^-1^ in ASW samples to 9,700 x 10^3^ copies L^-1^ in WW and CDW samples, then decreased to 2,400 x 10^3^ copies L^-1^ in SLOPE water samples. As a consequence of these distributions, the contribution of Thaumarchaeota to prokaryotes increased with depth, from a mean of 0.2% in ASW samples to a mean of 26% in SLOPE water samples.

Mean concentrations of *amoA* genes (WCA+WCB, 22) were 214 and 4,040 x 10^3^ copies L^-1^ in WW and CDW samples, respectively (Supplemental Table 4). These values are significantly lower than concentrations we measured in samples from the same water masses in 2011 (LMG1101, 23) using the Wuchter et al. (24) primer set: 4,100 and 12,500 x 10^3^ copies L^-1^, *p*<0.0001 and *p*=0.0002, respectively). We also found that the mean of the ratios of *amoA*/*rrs* genes in a given sample were lower on LMG1801 than LMG1101: 0.02 versus 1.7 (*p* <0.0001) and 0.46 versus 1.6 (*p* = 0.0005) for WW and CDW samples, respectively. The same *rrs* primers (25) were used in both studies, yielding much smaller, though statistically significant (*p*<0.0001), differences in *rrs* abundances between cruises: 9,700 versus 2,900 and 9,600 versus 16,000 x 10^3^ copies L^-1^ for WW and CDW samples collected on LMG1801 versus LMG1101. While some of the difference in *amoA* abundance between cruises may be attributed to interannual variability in the actual abundance or composition of AOA populations at the study site, it is more likely that it reflects amplification bias of the Mosier and Francis (22) versus Wuchter et al. (24) primers in our samples. Differences by water mass in the ratio of *amoA*:*rrs* in samples from LMG1801 suggest that *amoA* abundance is underestimated to a greater extent in WW populations, dominated by Shallow Water Clade A AOA, compared to CDW samples, dominated by Deep Water Clade B AOA (23).

*Nitrospina*, a dominant clade of nitrite oxidizers in the sea, may contribute to urease activity (26-28) and thus the production of NH_4_^+^ from urea. We detected *Nitrospina rrs* (29) throughout the water column (Supplemental Table 1) with greatest mean abundances in the WW and CDW water masses (675 and 583 x 10^3^ copies L^-1^, respectively (Supplemental Table 4), which were not significantly different (*p* = 0.095, Supplemental Table 6). The abundances of *Nitrospina rrs* in ASW and SLOPE water masses were lower and they were not significantly different from each other (mean abundances of 94 versus 180 x 10^3^ copies L^-1^, respectively, *p* = 0.50, Supplemental Tables 4 and 6).

Thaumarchaeota *ureC* genes (12) were also distributed throughout the water column, with greatest mean abundance (1,200 x 10^3^ copies L^-1^, Supplemental Table 4) in the CDW water mass. The distribution of Thaumarchaeota *ureC* was similar to that of Thaumarchaeota *rrs* and *Nitrospina rrs*, with lower concentrations in the ASW and SLOPE water masses (32 and 50 x 10^3^ copies L^-1^, respectively, Supplemental Table 4). Mean ratios of Thaumarchaeota *ureC*/*rrs* were 0.15 for samples from the ASW, 0.05 for the WW, 0.13 for the CDW, 0.02 for the SLOPE and 0.09 over all depths. Kruskal-Wallis ranks tests demonstrated that the ratios differed by water mass (Supplemental Table 6) and revealed that the median ratio for CDW samples was significantly greater (*p*<0.0001) than ratios for WW and SLOPE data, but that ratios for the other pairwise comparisons were not significantly different (*p*>0.01).

## DISCUSSION

### Response of rates to substrate amendments

Detection of N oxidation rates may require amendments of ^15^N-labeled substrates that significantly increase the concentration of total (labeled plus unlabeled) substrate in samples. Further, environmental concentrations of NH_4_^+^ and urea may fluctuate *in situ* depending on localized coupling between regeneration and uptake or oxidation, subjecting nitrifiers to short-term, temporal variation in substrate concentrations (14, 30, 31) that may influence rates. Elevated substrate concentration may influence nitrite production via enzyme kinetics (32, 33) or by increasing production of toxic by-products (34).

Inhibition of AO and UO in response to elevated substrate concentrations has been observed in other studies, but the phenomenon does not appear to have been fully assimilated into the conceptual model of Thaumarchaeota ecophysiology. AO and UO rates measured in samples from the 1% light level (51 m) during a period of active upwelling (March 2015) at the SPOT station off southern California (15) decreased in response to ^15^N amendments to samples with ambient NH_4_^+^ and urea-N concentrations of 10 nM and 190 nM (enrichment factors of 150-2,500% and 8-130%, respectively; Figure 5 in Laperriere et al, 15). Although not discussed in their paper, Shiozaki et al. (17) found that urea amendments of 1,560 nM (mean enrichment factor: 2,312%) inhibited UO rates 50 to 77% relative to rates measured with 31 nM amendments (mean enrichment factor: 145%) in 3 samples from the 0.1% light level in the Beaufort Sea (calculated from their Supplemental Dataset 1). They did not test the effect of NH_4_^+^ amendments on AO rates on this cruise; however, Shiozaki et al. (31) performed similar experiments with ^15^NH_4_^+^ amendments ranging from 31 to 1,560 nM (mean enrichment factors: 208 and 5,540%) using samples from the 0.1% light level at 15 stations on a meridional transect of the North Pacific. These experiments, reported in their Figure 4a and Supplemental Table 1, showed no clear response of AO to amendments: AO rates increased in 6 and decreased in 7 samples where AO rates were >LD. The mean change of AO rates with amendments of 1,560 nM versus 31 nM was 105% (range 44-273%). The smallest ^15^NH_4_^+^ amendments used in this study (31 nM) represent larger enrichment factors (194% to infinity for 12 samples from the North Pacific Gyre where ambient [NH_4_^+^] was undetectable in 3 samples, and 101-105% for 3 samples from the Bering Sea with high [NH_4_^+^]) than the 6 nM amendments used in our experiments (range 100-140% for both substrates).

**Figure 3.**
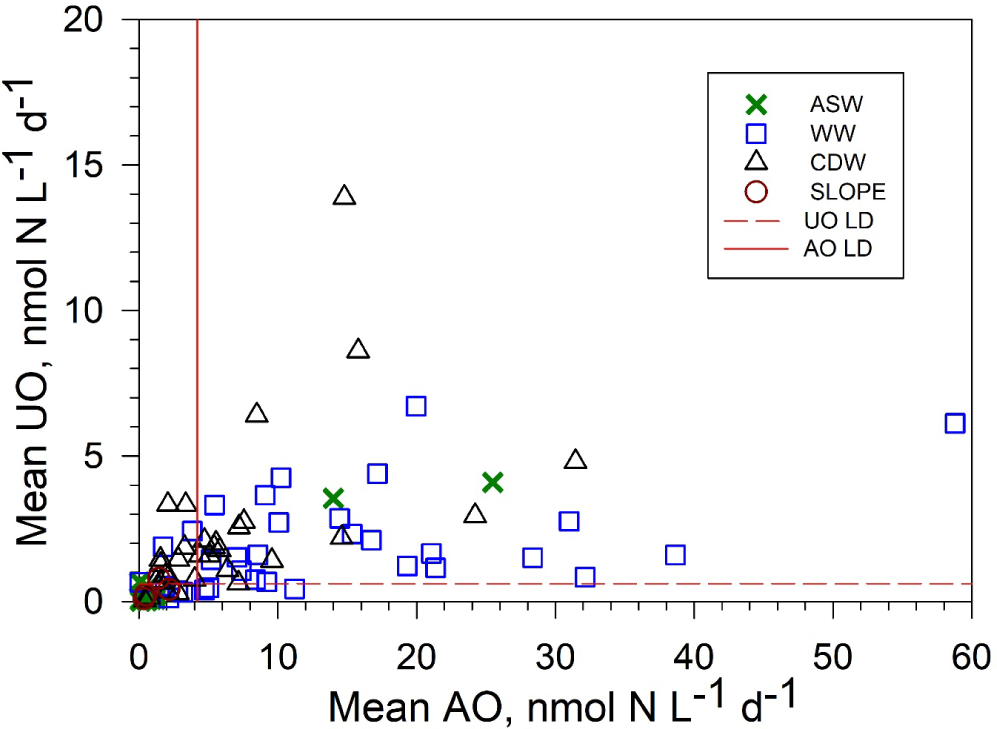
Oxidation rates of urea-N versus NH_4_^+^-N. Data points are means of duplicate UO and AO rates measured for a given sample. Red horizontal and vertical lines indicate the limits of detection estimated for these measurements. Outliers (WW: 114, 2.5 and 101, 3.8; CDW: 17.8, 117) have been omitted from the plot. ASW samples are shown as **x**, WW samples are shown as **□**, CDW samples are shown as Δ, and SLOPE samples are shown as ○.

**Figure 4.**
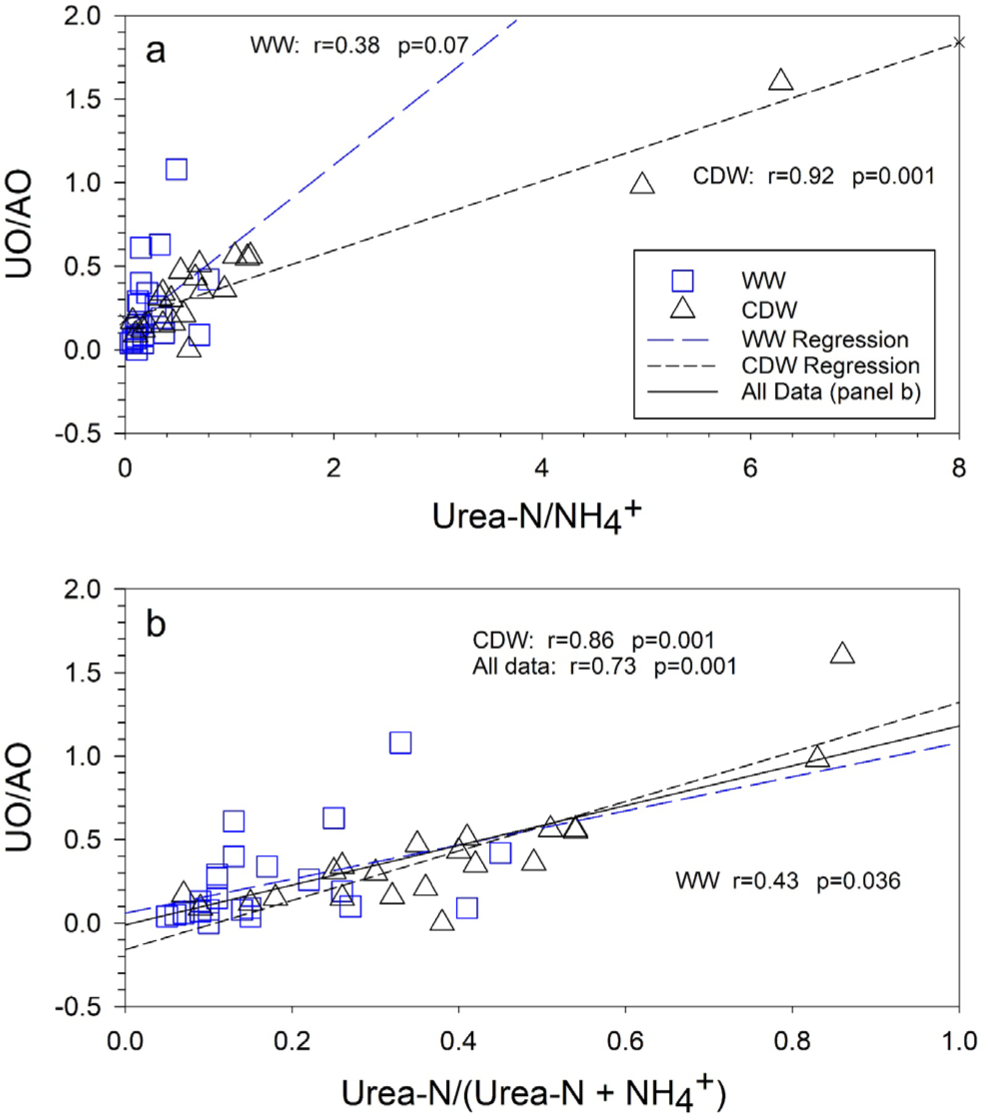
Ratio of the oxidation rates of urea-N to NH_4_^+^-N (UO/AO) versus: a) the ratio of [urea-N] to [NH_4_^+^-N] measured in the same sample; and b) the contribution of urea-N to oxidizable N ([urea-N]/([urea-N] + [NH_4_^+^-N])). UO and AO are means of duplicate rate measurements made for a particular sample (station and depth). Model 2 OLS regression lines, correlation coefficients and *p*-values for the correlation are shown. The heavy regression line in panel b) is for all data (WW + CDW combined). Symbols as in Figure 3.

A mechanism that might explain the response of CDW AOA to substrate amendments is sensitivity to reactive oxygen species (ROS) or reactive nitrogen species (RNS) produced during ammonia oxidation, particularly under conditions of elevated substrate concentrations (34). AOA are known to be inhibited by ROS species, including HOOH (34, 35), and previous work in our study area (36) verifies that these AOA populations are no exception. We hypothesize that ROS produced in response to elevated substrate concentrations caused by our amendments can reach toxic levels intracellularly or in the immediate vicinity of the cells, inhibiting further oxidation of N supplied as NH_4_^+^ or urea. The greater inhibition of CDW populations by NH_4_^+^ vs urea may be due to the slower rate at which N from urea versus NH_4_^+^ is oxidized, and thus ROS/RNS is produced, in these samples (21.2 vs 1.6 nmol N L^-1^ d^-1^ for AO vs UO, respectively, for all WW samples; 7.9 vs 2.5 (outlier excluded), in all CDW samples, Supplemental Table 4).

It is also likely that sensitivity to, or production of, ROS/RNS varies among AOA clades (34). Gene ratios in samples collected on LMG1801 (Supplemental Figure 5), as well as more rigorous analyses performed previously (23, 37, 38), demonstrate that WW and CDW Thaumarchaeota populations are phylogenetically distinct. This difference may influence the rates at which they produce, or detoxify, ROS/RNS.

Alternatively, elevated [NH_4_^+^] may result in a shift in the N oxidation pathway resulting in an increase in the ratio of N_2_O:NO_2_^-^ produced, as reported by Frey et al. (33). Our experimental protocol would not have captured ^15^N_2_O or other gaseous intermediates, potentially underestimating N oxidation rates.

Our observations (Figure 1) and those of others cited above suggest that amendments that increase substrate concentrations significantly can affect the rates measured, and not as expected from simple Michaelis-Menten enzyme kinetic considerations. Over our entire data set of >200 measurements, amendments of 44 nM ^15^NH_4_^+^ resulted in enrichment factors of 124% ± 29% (mean ± SD); 112 ± 12% in WW samples and 146 ± 28% in CDW samples. ^15^N-urea amendments (47 nM) resulted in enrichment factors of 274 ± 256%: 319 ± 382% and 290 ± 190% in samples from the WW and CDW, respectively. Assuming that AOA populations in our samples responded to substrate enrichments similarly to those in our experiments, and that 6 nM represents a true “tracer” amendment, the AO and UO rates we measured in WW samples may overestimate *in situ* rates by 184 and 110%, respectively, while those for CDW samples may be 25% and 77% of the *in situ* rates, on average. Differential inhibition may account for the differences in mean rates (Figure 2, Supplemental Table 4) and in mean cell-specific rates (31 vs 17 fmol N cell^-1^ d^-1^, Mann-Whitney 2-tail *p* = 0.0063) between WW vs CDW samples. We have not corrected the data reported in Supplemental Table 1 for these differences.

Rate measurements made in open ocean samples where [NH_4_^+^] and [urea] are in the low nM range typically use substrate amendments that range from 30 to 50 nM. Measured rates are thus likely to have been affected by the change in substrate concentration due to the tracer amendment. The data suggest that the effect is very nonlinear (Figure 1; Figure 5 in Laperriere et al, 15; Figure 4a in Shiozaki et al, 17; Shiozaki et al, 31; Kim et al, 34). Comparisons between rates measured with 30-50 nM additions and rates measured with much higher substrate additions may show little response (e.g. Shiozaki et al, 31) because the threshold for response is lower than 30-50 nM. Table 1 compares rates we measured with 44 or 47 nM amendments with those measured with 440 or 470 nM amendments, by water mass. Although the differences are not statistically significant, responses of rates to 440 vs 44 nM amendments are damped relative to 44 vs 6 nM amendments, especially for AO. Finally, our data suggest that, compared to WW populations, CDW AOA are poorly adapted to fluctuations in substrate concentrations that might arise from uncoupling between production and consumption of NH_4_^+^ (14, 30, 31), or patchiness (39-41). This may be a defining characteristic of the ecophysiology of epipelagic versus mesopelagic AOA.

**Table 1.** Comparison of the response of AO and UO rates to amendments of 44 vs 6 or 440 vs 44 nM amendments. *P*-values are for 2-tailed *t*-tests.

|  | Ratio of rates with 44 vs 6 nM amendments |  | Ratio of rates with 440 vs 44 nM amendments |  | *p* |
| --- | --- | --- | --- | --- | --- |
|  | Mean | n | Mean | n |  |
| AO WW | 1.84 | 2 | 1.01 | 3 | 0.062 |
| AO CDW | 0.25 | 3 | 0.42 | 4 | 0.069 |
| UO WW | 1.28 | 3 | 0.84 | 3 | 0.101 |
| UO CDW | 0.77 | 3 | 0.72 | 3 | 0.765 |

### Relationships among variables

We compared the distribution of AO and UO to the distribution of relevant marker genes and environmental variables (Supplemental Figures 6 and 7, Supplemental Table 7). Rates that were <LD were set to 0 for this analysis (21), although using all data yielded essentially the same results. We found statistically significant correlations between the abundance of *Nitrospina rrs* genes and AO or UO rates (AO all data: r^2^=0.43, *p*=0.001; UO all data: r^2^=0.21, *p*=0.004). The relationships were stronger for WW samples than for CDW samples (Supplemental Table 7). The “reciprocal feeding” model (42) for the role of *Nitrospina* in ammonia oxidation predicts a positive relationship between *Nitrospina* abundance and AO. While urease activity associated with *Nitrospina* may be an explanation for the correlations we observed, the correlation could also be based on other factors, such as urea supply or the rate of nitrite production in a sample by combined AO + UO.

AO rates were significantly positively correlated with [NH_4_^+^] in CDW samples and weakly correlated with the abundances of Thaumarchaeota *rrs* and with *Nitrospina rrs* genes (Supplementary Figure 6, Supplemental Table 7: for all samples, Thaumarchaeota *rrs* genes r=0.26, *p*=0.002). UO rates were significantly positively correlated with both [NH_4_^+^] and [urea] in CDW samples.

The abundance of Thaumarchaeota *ureC* genes was significantly correlated with the abundance of Thaumarchaeota *rrs* genes (Supplemental Table 7). We found no significant correlations between the abundance of Thaumarchaeota *ureC* genes and either [NH_4_^+^] or [urea], or with the ratio ([urea-N]/[NH_4_^+^]), or with the ratio ([urea-N]/([urea-N + NH_4_^+^])), in any of the water masses we sampled (Supplemental Figure 5, panels g and h). The ratio of Thaumarchaeota *ureC* to Thaumarchaeota *rrs* genes was greatest in CDW samples (regression slope = 0.13, mean of the ratio of *ureC*/*rrs* for data from the same sample = 0.13) and distinct from the ratio in WW samples (regression slope 0.03, mean ratio of *ureC*/*rrs* = 0.05; Supplemental Tables 4 and 7).

These *ureC*/*rrs* ratios are lower than those reported by Alonso-Sáez et al. (12): 0.09 vs 0.76 for all of their data, 0.09 vs 0.51 for their data with an outlier removed, (*p*<0.0001 in both cases), and did not increase with depth (model 2 r = -0.13, *p*=0.08). We examined their data, reported in their supplemental tables S4 and S5. The relationship between Thaumarchaeota *ureC*/*rrs* and depth was strongly influenced by the value of one outlier that was based on a *ureC* analysis with a very high standard deviation (mean±SD = 21.95±10.09). The correlation between the ratio of Thaumarchaeota *ureC*/*rrs* and depth was not statistically significant, regardless of whether the outlier is included (r^2^=0.03, *p*=0.15), or not (r^2^=0.03, *p*=0.16). Within the CDW data set that was the basis for the conclusion that Thaumarchaeota *ureC*/*rrs* ratios increase with depth, the mean Thaumarchaeota *ureC*/*rrs* ratio was 2.67, but without the outlier it was 1.04. The ratio of Thaumarchaeota *ureC*/*rrs* does not appear to be a good predictor of the contribution of urea to nitrification, and there seems to be little change with depth in the contribution of urea to nitrite production, at least in our study area.

The means of duplicate measurements of AO and UO were not correlated: r=0.12, *p*=0.24 for WW samples and r=0.13, *p*=0.14 for CDW samples (Figure 3). The mean ratio of UO/AO from the complete data set (n=43 for samples where rates of both replicate analyses were >LD) was 0.39 with a range of 0.02-6.6 (Supplemental Table 7). The ratio of 6.6 was from one sample with an unusually high urea concentration. The mean ratio is 0.25, with a range of 0.02-0.94, when this outlier is excluded from the calculation. Ratios of UO/AO measured in the WW water mass averaged 0.18 while those in samples from the CDW water mass averaged 0.64 (0.33 with the outlier excluded). These values are significantly different (Mann-Whitney ranks tests, *p* = 0.013, CDW outlier removed). Wan et al. (14) also found that UO/AO increased with depth based on samples from 4 depths at 4 stations in the north Pacific. This trend might be an artifact of greater inhibition of AO than UO in CDW (mesopelagic) samples due to ^15^N amendments.

### Contribution of urea-N to nitrification

We explored the relationships between rates and variables, or combinations of variables, related to AO and UO to determine if they could be used to predict activity. We found no statistically significant relationships between UO and [NH_4_^+^] or [urea] when all samples were considered together, or in the subset of WW samples (Supplemental Table 7). We found that UO was positively correlated with both [NH_4_^+^] and [urea] in CDW samples (Supplemental Table 7), indicating that UO was not inhibited by [NH_4_^+^], in contrast to experiments with Chukchi Sea populations (17).

UO correlated significantly with the contribution of urea-N to oxidizable N, approximated as ([urea-N]/([urea-N]+[NH_4_^+^])), when all samples were considered together (*p*=0.003), but the correlation was weak (r=0.23) and was not significant when considered by water mass (Supplemental Table 7). The ratio of rates (UO/AO) was predicted by the ratio of [urea-N] to [NH_4_^+^] (Figure 4, panel a); however, this relationship was not consistent between water masses (WW slope = 0.49, r=0.38, CDW slope = 0.21, r=0.92). The best predictor of UO/AO in a sample was ([urea-N]/([urea-N]+[NH_4_^+^])) in the same sample (Figure 4, panel b). With the caveat that the number of samples from each water mass with data allowing the calculation of both parameters was small, we found that the strength of this relationship differed between water masses: r=0.43 for WW samples, but r=0.86 for samples from the CDW and r=0.73 for the combined WW+CDW data set. The slopes of the regressions (Figure 4b: 1.01, 1.48 and 1.19 for WW, CDW and All Data, respectively) were not significantly different (p<0.05). However, neither of these parameters was a good predictor of the absolute rate of UO. And, while we found a strong relationship between UO/AO and ([urea-N]/([urea-N]+[NH_4_^+^])) in our study area, this relationship does not hold for data from studies of other locations (Gulf of Mexico, 16; Arctic Ocean, 17), where all required variables are available for this analysis.

Supplemental Table 8 compares data from LMG1801 with ratios of the oxidation rates of N supplied as urea versus NH_4_^+^ (UO/AO) calculated from data in other studies. Data from LMG1801 indicate that the contribution of urea-N to nitrite production on the continental shelf west of the Antarctic Peninsula was ∼25% of that produced by AO, and that its contribution became relatively more important as the contribution of urea to oxidizable N increases. Values at other locations range from very small contributions of urea to nitrite production (Gulf of Mexico) to urea supplying most of the N oxidized to nitrite (LMG1101 WW, deep water at the SPOT time series station, Bering/Chukchi Seas). We found no relationship between UO/AO and measures of the relative availability of urea-N in the other data sets we examined, including our data from the South Atlantic Bight (43). SPOT data (15) suggest an increase with depth in the contribution of urea to nitrification, as do data presented by Wan et al. (14), and as we found on LMG1801; however, data reported by Shiozaki et al, (17) have the opposite trend (contribution of UO decreases with depth). These data demonstrate that the contribution of urea to nitrification in the open ocean can be significant, but it appears to be highly variable and the data do not support the general conclusion that the contribution of urea-N to nitrite production is enhanced in Antarctic coastal (polar) waters relative to sites at lower latitudes (12).

## CONCLUSIONS

The response of N oxidation rates to substrate amendments was complex, with measured rates increasing slightly with increases in total substrate concentration for WW samples, but strongly inhibited in CDW samples. Inhibition may have been caused by increased production of reactive oxygen or nitrogen species accompanying oxidation of NH_4_^+^ or urea-N, or by shifts in the end-product from nitrite to N2O, as total substrate concentration increased. This may be a general problem for rate measurements made in samples from the mesopelagic zone of the open ocean and suggests that mesopelagic Thaumarchaeota populations are not well-adapted to short-term fluctuations in substrate concentration.

Urea-N contributed significantly to the production of nitrite in samples from the continental shelf and slope west of the Antarctic Peninsula. Oxidation rates of urea-N were 25%, on average, of the oxidation rates of NH_4_^+^, similar to the contribution of urea to nitrite production in Georgia coastal waters (43) and in contrast to a greater contribution of urea to nitrite production in polar waters suggested by others (12). Oxidation rates of urea-N were not correlated with the ratio of Thaumarchaeota *ureC*/16S rRNA, nor with [NH_4_^+^], [urea] or rates of NH_4_^+^ oxidation. Oxidation of urea-N was not inhibited by elevated NH_4_^+^ concentrations.

## MATERIALS AND METHODS

A more detailed description of sample collection, processing and analysis is presented in the Supplemental Material linked to this article.

### Sample Collection

We sampled the continental shelf and slope west of the Antarctic Peninsula (Supplemental Figure 1) during the austral summer of 2018 (ARV Laurence M Gould cruise LMG1801, PAL-LTER cruise 26, DOI: 10.7284/907858). Sampling focused on 3 or 4 depths at each station, chosen to represent Antarctic Surface Water (ASW, samples from 10 or 15 m), Winter Water (WW, samples from 35 to 100 m, targeting the water column temperature minimum), Circumpolar Deep Water (CDW, samples from 175 to 1,000 m depth) and slope water (SLOPE, samples from 2,500 to 3,048 m depth, generally ∼10 m above the bottom at stations on the slope or over basins on the shelf). Water was collected in Niskin bottles (General Oceanics Inc., Miami, FL, USA). Samples for DNA and nutrient analyses were drained into opaque 2 L HDPE plastic bottles. Water for incubations was drained into aged, acid-washed, sample-rinsed polycarbonate bottles (Nalge) that were kept in cardboard boxes to minimize exposure to light.

DNA samples were filtered under pressure through 0.22 μm pore size Sterivex filters (EMD Millipore, Billerica, MA, USA). Residual seawater was expelled, then lysis buffer (0.75 M sucrose, 40 mM EDTA, 50 mM Tris, pH 8.3) was added to the filter capsule, which was capped, frozen, then stored at -80 °C. Samples of the Sterivex filtrate were frozen at -80 °C for subsequent chemical analyses. One set of filtrate samples was stored briefly at 4 °C, then used for onboard determination of ammonium concentration by the *o*-phthaldialdehyde method (44). Urea was determined manually from frozen samples by the diacetyl monoxime method (45, 46).

### Gene abundance

DNA was recovered from Sterivex filters using a lysozyme and proteinase K digestion, followed by phenol-chloroform extraction (47). Archaea and Bacteria genes in the extracts were quantified by PCR (qPCR). The primers and probes used, PCR reaction conditions and our estimates of the precision of the measurements are given in Supplemental Table 9.

### Nitrogen oxidation rates

AO and UO were measured using ^15^N-labeled substrates. Substrates were added to samples within ∼1 hr of collection to yield ∼44 nM of ^15^NH_4_^+^ (32, 48, 49) or ∼47 nM of urea (94 nM of urea-^15^N). These amendments increased substrate concentrations in the samples ((([^15^N amendment] + [ambient])/([ambient]))*100) by an average of 125%, range: 101-202% and 102-1,800% for NH_4_^+^ and urea, respectively. Labeled substrates were added to duplicate bottles that were incubated in the dark for ∼48 hr. Incubation temperature averaged 0.23 °C with a standard deviation of 0.71 °C. Incubations were terminated by decanting ∼40 mL subsamples into plastic tubes that were immediately frozen at -80 °C.

We ran experiments with samples from 2 depths at 3 stations to verify that ^15^N oxidation rates did not change significantly during incubations (Supplemental Figure 8), to assess the effect of substrate amendments on measured rates (Figure 1, Supplemental Figure 2), and to assess the effect of incubation temperature on measured rates (Supplemental Figure 3). The characteristics of the samples used in these experiments compared favorably (Supplemental Table 4, *t*-test, *p*>0.01) with mean conditions over all samples from the same water mass, with few exceptions: the concentrations *ureC* and WCB *amoA* genes and T in the WW sample from Station 600.040B; AO in the WW sample from Station 149.-050; and the concentrations of NH_4_^+^ and Bacteria *rrs* in the WW sample from Station 200.000, which were all significantly greater than water mass means (Supplemental Table 4). Rates calculated from single-point determinations, (end-points of samples from the survey, from experiments, or the 48 hr points from time courses), agreed well with rates estimated from the slopes of regressions of time course data (Supplemental Table 10). Rates estimated from slopes were generally lower than rates calculated from end-point determinations, which assume intercepts of 0, while intercepts of regressions ranged from -0.25 to 1.41 nmol L^-1^.

### ^15^N in nitrite plus nitrate

The ^15^N content of NO2-plus NO3-(^15^NO_x_) of our samples was measured using the ‘denitrifier method’ (50) with *Pseudomonas aureofaciens* as described previously (49). The N_2_O produced was analyzed using a Gas Bench II coupled to a Finnegan MAT 252 mass spectrometer (51, 52).

### Rate calculations

Our rate measurements are based on the production of ^15^NO_x_ from ^15^N labeled substrates. We calculated oxidation rates by comparing δ^15^N values of the NO_x_ pool at the ends of the incubations with values in unamended samples (“natural abundance”), as described previously (49). We assumed that the δ^15^N value of naturally occurring ammonium and urea is the same as that of N_2_ in air. Chemical data needed for rate calculations were not available for some samples (see Supplemental Table 1), so we substituted water mass averages (Supplemental Table 4) determined from other samples taken on the cruise. Samples with low or no activity sometimes yielded negative rates because the δ^15^NO_x_ “natural abundance” value determined for that sample was greater than the δ^15^NO_x_ value determined for the amended treatment sampled at the end of the incubation. These values were set to 0 (21) for statistical analyses. Note that the rates we report are for N oxidized, regardless of whether it was supplied as NH_4_^+^ or urea.

### Precision and accuracy

Analytical uncertainty of δ^15^N measurements ranged from 0.36‰ to 0.56‰. Accuracy was 0.42‰ (at-% ^15^N = 0.00019, n = 56). The precision of nitrite+nitrate analyses run by PAL-LTER personnel was reported to be 100 nM. We determined the precision of ammonium and urea analyses as the mean standard deviation of replicate (2 or 3) analyses of a given sample. They are: ammonium, 65 nM; urea, 10 nM. We ran 10,000 Monte Carlo simulations using cruise means of these variables and their precisions to estimate the precision of the resulting rate measurements. These are: 2.3 nmol N L^-1^ d^-1^ for AO and 0.31 nmol N L^-1^ d^-1^ for UO; for relative standard deviations (RSD; ((standard deviation/mean) x 100)) of 15% and 11%, respectively, of calculated rates. The limit of detection for a measurement was set at 1.96 times the precision of the measurement.

### Statistical analyses

Rates that were below the limits of detection as established above, were assigned values of 0 (21). We tested for spatial gradients in the distributions of variables across the study area within a water mass (Supplemental Figure 4) by grouping stations by location (northeast versus southwest, inshore versus offshore), as shown in Supplemental Figure 1. Assignments of individual stations to these groups are given in Supplemental Table 1. We used Mann-Whitney ranks tests to determine if variables were distributed uniformly across the study area within a water mass, and Kruskal-Wallis ranks tests of the significance of differences between the 4 water masses sampled. Variables that were not uniformly distributed among water masses (most of them) were analyzed further using *post hoc* Dunn tests, with *p*-values adjusted for false discovery rate using the Benjamini-Hochberg correction, to identify sets that differed significantly at *p*<0.01. Pearson product moment regressions run in VassarStats (http://vassarstats.net/, ^©^R. Lowry) were used to obtain slopes of time courses. We used model 2 ordinary least square regressions run in R (53) to test for correlations between variables.

### Data archives

The data we collected on LMG1801 are archived by the Biological and Chemical Oceanography Data Management Office (BCO-DMO) under project acronym “Oxidation of Urea N,” doi:10.26008/1912/bco-dmo.840629.2, https://www.bco-dmo.org/dataset/840629/data.The data used in the analyses presented here are reported in Supplemental Table 1, with summaries by water mass given in Supplemental Table 4.

## ACKNOWLEDGMENTS

We thank the officers and crew of the ARSV Laurence M Gould and staff of Raytheon Polar Services Company, especially Diane Hutt, for their support during cruise LMG1801, and personnel affiliated with the Palmer LTER (funded through Grant NSF PLR 1440435) for additional support on LMG1801 and for subsequent access to project data. We would also like to thank S. Rauch at BCO-DMO for her assistance in archiving the data from this project and T. Hastings for just being there. This work was supported by the US National Science Foundation through grants OPP 1643466, (to JTH) and OPP 1643354 (to BNP). This is SOEST contribution number XXXX.

## AUTHOR CONTRIBUTIONS

JTH and BNP designed the research; JTH, BNP and HD conducted the sampling program; JTH, JD, AO-O, NJW, TA and BNP contributed to sample analysis; JTH and BNP analyzed the data, JTH wrote the paper with input from the coauthors.

---

**Supplemental Figure 1.** Chart of the study area. The orange double line separates stations assigned to the NE vs SW groups. Symbols for nearshore stations are green squares, symbols for offshore stations are blue circles. Stations used to validate our experimental protocols are indicated by an X. Line numbers correspond to the PAL LTER grid numbering system (https://pallter.marine.rutgers.edu/). Base map courtesy LTER Network Office (https://lternet.edu/).

**Supplemental Figure 2.** Oxidation rates of N supplied as NH_4_^+^ (AO) or urea (UO) as functions of ^15^N-labeled substrate amendments (as nmol L^-1^ of the substrate, not of N in the case of urea). Solid bars are WW samples, cross-hatched bars are CDW samples. Sample depths are given in Supplemental Table 1.

**Supplemental Figure 3.** Response of AO rates to incubation temperature. Points from duplicate rate measurements overlap in some cases. Primary data (panel a) were transformed as the square root of the data normalized against the highest rate recorded (panel b; 54, 55).

**Supplemental Figure 4.** Distribution of variables related to the oxidation of NH_4_^+^ or urea-N across the study area, by water mass. The data for a given variable from a given water mass were tested (see Supplemental Table 5) for random distribution between pairs of geographic groups as indicated in Supplemental Figure 1. See Supplemental Table 1 for assignments of individual stations to groups. The areas of the circles on each plot are scaled to values of the variable, with a key given at position: (latitude, longitude) -62, -76 on each panel. The key also shows the locations of all samples taken from a given water mass. Measurements that were below the limits of detection(LD) have been set to 0 and thus are not shown on the plots. Sample temperatures were re-scaled to values >0 °C by adding 2 °C to all measured values. Base map courtesy LTER Network Office (https://lternet.edu/). Columns (left to right): 1, abundance of Bacteria 16S rRNA genes (*rrs*, 10^9^ copies L^-1^, LD=0.01); 2, Thaumarchaeota 16S rRNA genes (*rrs*, 10^3^ copies L^-1^, LD=3.9); 3, Thaumarchaeota ammonia monooxygenase genes (*amoA*, 10^3^ copies L^-1^, LD=2.0); 4, the α subunit of Thaumarchaeota urease (*ureC*, 10^3^ copies L^-1^, LD=15.7); 5, *Nitrospina* 16S rRNA genes (*rrs*, 10^3^ copies L^-1^, LD=3.9); 6, oxidation rate of NH_4_^+^ N (AO, nmol N L^-1^ d^-1^, LD=4.3); 7, oxidation rate of urea-N (UO, nmol N L^-1^ d^-1^, LD=0.61); 8, sample temperature (°C + 2); 9, sample salinity (PSU).

**Supplemental Figure 5.** Biplots of gene abundances by water mass. ASW omitted because of minimal data. a) Thaumarchaeota *amoA* vs Thaumarchaeota *rrs*, b) Thaumarchaeota *ureC* vs Thaumarchaeota *rrs*, c) Thaumarchaeota *ureC* vs Thaumarchaeota *amoA*, d) Thaumarchaeota *ureC* vs Nitrospina *rrs*, e) Thaumarchaeota *ureC* vs [urea], f) Thaumarchaeota *ureC* vs [NH_4_^+^], g) Thaumarchaeota *ureC* vs ([urea]/[NH_4_^+^]), h) Thaumarchaeota *ureC* vs [urea-N]/([urea-N]+[NH_4_^+^]). Slopes, coefficients of determination and *p*-values of the correlation (“NS” = *p*>0.05) are from model II ordinary least squares regressions. Trend lines are shown for significant (*p*<0.05) regressions. The legend in panel a) shows line styles used for each water mass. Samples from the WW water mass are shown as **□**, samples from CDW are shown as Δ, and samples from SLOPE water are shown as X. Outliers have been omitted from some of the plots (see panels) to improve the resolution of points near the origins.

**Supplemental Figure 6.** Oxidation rates of N supplied as NH_4_^+^ versus values of selected environmental variables measured in the same sample. a) Thaumarchaeota *rrs*, b) Thaumarchaeota *amoA*, c) Thaumarchaeota *ureC*, d) Nitrospina *rrs*, e) [NH_4_^+^], f) [urea]. Samples from the WW are shown as **□**, samples from the CDW are shown as Δ, and samples of SLOPE water are shown as X. Red horizontal lines indicate the limits of detection for rate measurements. The significance of model 2 regressions of subsets of the data are given in Supplemental Table 7.

**Supplemental Figure 7.** Oxidation rates of N supplied as urea (UO) versus values of selected environmental variables measured in the same sample. a) Thaumarchaeota *rrs*, b) Thaumarchaeota *amoA*, c) Thaumarchaeota *ureC*, d) Nitrospina *rrs*, e) [NH_4_^+^], f) [urea], g) ratio ([urea-N]/[NH_4_^+^]), h) urea availability ([urea-N]/([urea-N] + [NH_4_^+^])). The significance of model 2 regressions of subsets of the data are given in Supplemental Table 7. Symbols as in Supplemental Figure 6. Some points have been omitted from the plots (see panels) to improve the resolution of points near the origins.

**Supplemental Figure 8.** Time courses of the production of ^15^NO_x_ from ^15^N-labeled NH_4_^+^ and urea. Samples were collected at the stations and depths indicated, replicate 250 mL bottles were amended with 44 or 47 nM ^15^N-labeled NH_4_^+^ or urea, respectively, then incubated in the same incubator as survey measurements. Duplicate bottles were removed at the times shown, 40 mL was decanted from each bottle into a centrifuge tube and frozen at -80 °C until they could be analyzed for ^15^NO_x_ content. Time course data were analyzed to determine the slope of the Pearson product moment regressions shown as dashed lines if r^2^>0.5.

**Supplemental Table 1.** Data collected on cruise LMG1801. The two rows labeled “Measurement Precision” and “Limit of Detection” provide estimates of those values for the data in the columns below the entries. See text for details. Column headings give measurement names and units and are generally self-explanatory. Cells in the “Experimental Replicate” column containing the text “48 hr”, “44 nM” and “T=0” are from experiments to verify our protocols (respectively: time courses, concentration dependence, and temperature dependence). Replicates from survey measurements are labeled “A” and “B”. Environmental and qPCR data for a given sample are listed with the “A” replicate of survey measurements, though they also apply to the “B” replicate. Blank cells indicate no data. Outliers enclosed in parentheses have been excluded from calculations of descriptive statistics (presented in Supplemental Table 4) for the water mass in which they occur. Shading indicates water mass designation (ASW 0-34 m; WW 35-100 m; CDW 175-1,000 m; SLOPE 2,500-3,048 m).

**Supplemental Table 2.** Comparisons of changes in the oxidation rates of N supplied as NH_4_^+^ or urea in response to substrate amendments. Rates shown in Supplemental Figure 2 were normalized as percentages of the highest rate in a set (substrate, station, water mass, amendments being compared; e.g. NH_4_^+^, Station 600.180, WW, 6 vs 44 nM). The mean normalized scores (e.g. NH_4_^+^, all stations, WW, all 6 nM amendments) were calculated and are reported in the top 3 rows of each section. The *p*-values reported are for 2-tailed *t*-tests of the significance of the difference between the normalized scores for the two amendments being compared (n1 and n2 independent samples, unequal sample variance). Values of *p*<0.01 are shown in **BOLD**.

**Supplemental Table 3.** Q_10_ and T_min_ values calculated from data in Supplemental Figure 3. T_min_ values calculated as per (54).

**Supplemental Table 4.** Descriptive statistics of water mass properties and comparison to values from samples used to test experimental protocols. Columns at right give the means of duplicate rate measurements made for that sample (station and depth, Supplemental Table 1). Means that were less than the limit of detection (<LD) were excluded from further calculations. AO = oxidation of ammonia N, UO = oxidation of urea-N. Rows at the bottom of the table compare values of variables and parameters from samples used in tests with the mean value from the same water mass. Values that are significantly different from the water mass mean at p<0.01 (mean ± (2.3263 * stdev.s)) are indicated in *BOLD RED ITALICS*. Blank cells indicate no data for that variable or parameter. Shading highlights water mass designations.

**Supplemental Table 5.** Results of Mann-Whitney ranks tests of the distribution of variables across the study area by sampling day and geographic location. Areal distributions of the data by water mass are shown in Supplemental Figure 4. The stations were assigned to subsets by sampling day (“Days 1-15” vs “Days 16-30”) and geographic region (“Northeast” vs “Southwest” or “Inshore” vs “Offshore,”), see Supplemental Table 1 for assignments of individual stations to groups. Data from a given water mass were tested to determine if their distribution between subsets was random (H_0_ is that there is no difference between subsets, rejected if *p*<0.01, highlighted in ***BOLD RED ITALICS***). Values given are the means of each subset followed by the probability that the distribution of values between subsets is random. One outlier from the CDW, offshore, urea data (2,060 nM) was excluded from calculations. We did not test the ASW or SLOPE data sets because most of the samples from those water masses were collected during the first half of the cruise (days 1-15). The SLOPE data sets are small (n≤16, including duplicate measurements of the same sample), there were too few measurements of the abundance of some genes in ASW samples, and too many values of ammonia and urea oxidation rates in the ASW water mass were below the limit of detection, thus assigned values of 0, for tests of spatial distributions within this water mass to be meaningful.

**Supplemental Table 6.** Results of Kruskal-Wallis ranks tests of the uniformity of the distribution of variables among water masses.

**Supplemental Table 7.** Summary of Model II, ordinary least squares regressions of variables related to the oxidation of N supplied as NH_4_^+^ or urea in samples collected on LMG1801. n is the number of observations, r is the correlation coefficient, *P* is the probability that the slope ≠ 0 and was derived from 999 bootstrap iterations. Rates <LD were assigned values of 0 for the analyses. Ratios used means of duplicate rates measured in a given sample where both rates are >LD. AO – oxidation of NH_4_^+^-N; UO – oxidation of urea-N.

**Supplemental Table 8.** Contribution of urea-N relative to NH_4_^+^ to nitrite production measured in other studies.

**Supplemental Table 9.** Primers and probes used in this study, qPCR cycling program, number of plates run, primer efficiencies and limits of detection.

**Supplemental Table 10.** Comparison of N oxidation rates from time courses of ^15^NO_x_ production with measurements from other experiments with the same sample. Rates were calculated from time courses as the slopes of Pearson product-moment regressions and are reported as “rate (r^2^, lower 99% CL-upper 99% CL).” “Rates from 48 hr points” are calculated from samples taken at ∼48 hours during time course incubations. “Rates from end-point determinations” are from incubations that were only sampled once after ∼48 hr of incubation. “Survey” samples were from the survey of nitrification rates across the study area. “44 nM” are from samples amended with 44 or 47 nM of ^15^N-labeled substrate as part of a study of the response of nitrifiers to higher or lower substrate concentrations. “Temp = 0” samples were part of a study to assess the effect of incubation temperature on rates. “Rep A” and “Rep B” indicate separate independent incubations (replicates). “dup” indicates samples for which ^15^NO_x_ analyses were replicated. “ASW,” Antarctic Surface Water, samples from 10-15 m; “WW,” Winter Water, samples from the temperature minimum between 35-100 m; “CDW,” Circumpolar Deep Water, 175-1000 m. Rates that are significantly different (99% CL) from rates determined by time course regressions are indicated by ***BOLD RED ITALICS***.

